# Predicting protein targets for drug-like compounds using transcriptomics

**DOI:** 10.1101/254367

**Authors:** Nicolas A. Pabon, Yan Xia, Samuel K. Estabrooks, Zhaofeng Ye, Amanda K. Herbrand, Evelyn Süß, Ricardo M. Biondi, Victoria A. Assimon, Jason E. Gestwicki, Jeffrey L. Brodsky, Carlos J. Camacho, Ziv Bar-Joseph

**Affiliations:** Department of Computational and Systems Biology, University of Pittsburgh, Pittsburgh, PA 15213; Machine Learning Department, School of Computer Science, Carnegie Mellon University, 15213; Department of Biological Sciences, University of Pittsburgh, Pittsburgh, PA 15260; School of Medicine, Tsinghua University, Beijing, China 100084; Department of Internal Medicine I, Universitätsklinikum Frankfurt, 60590 Frankfurt, Germany; Department of Pharmaceutical Chemistry, University of California, San Francisco, San Francisco, CA 94158

## Abstract

The development of an expanded chemical space for screening is an essential step in the challenge of identifying chemical probes for new, genomic-era protein targets. However, the difficulty of identifying targets for novel compounds leads to the prioritization of synthesis linked to known active scaffolds that bind familiar protein families, slowing the exploration of available chemical space. To change this paradigm, we validated a new pipeline capable of identifying compound-protein interactions even for compounds with no similarity to known drugs. Based on differential mRNA profiles from drug treatments and gene knockdowns across multiple cell types, we show that drugs cause gene regulatory network effects that correlate with those produced by silencing their target protein-coding gene. Applying supervised machine learning to exploit compound-knockdown signature correlations and enriching our predictions using an orthogonal structure-based screen, we achieved top-10/top-100 target prediction accuracies of 26%/41%, respectively, on a validation set 152 FDA-approved drugs and 3104 potential targets. We further predicted targets for 1680 compounds and validated a total of seven novel interactions with four difficult targets, including non-covalent modulators of HRAS and KRAS. We found that drug-target interactions manifest as gene expression correlations between drug treatment and both target gene knockdown and up/down-stream knockdowns. These correlations provide biologically relevant insight on the cell-level impact of disrupting protein interactions, highlighting the complex genetic phenotypes of drug treatments. Our pipeline can accelerate the identification and development of novel chemistries with potential to become drugs by screening for compound-target interactions in the full human interactome.

## Background

Most protein research still focuses on roughly 10% of proteins, and this bias has a profound effect on drug discovery, as exemplified by the popular kinase target [1-3]. The origin for this relatively limited exploration of the human interactome and the resulting lack of novel drugs for emerging ‘genomic-era’ targets has been traced back to the availability of small molecular weight probes for only a narrow set of familiar protein families [1]. To break this vicious circle, a new approach is needed that goes beyond known targets and old scaffolds, and that benefits from the vast amount of information we now have on gene expression, protein interactions, their structures and related diseases.

The current target-centric paradigm relies on high-throughput *in-vitro* screening of large compound libraries against a single protein [4]. This approach has been effective for kinases, GPCRs, and proteases, but has produced meager yields for new targets such as protein-protein interactions, which require chemotypes often not found in historical libraries [5, 6]. Moreover, these *in-vitro* biochemical screens often cannot provide any context regarding drug activity in the cell, multi-target effects, or toxicity [7, 8]. On the other hand, the goal of leveraging new chemistries would entail a compound-centric approach that would test compounds directly on thousands of potential targets. In principle, this is regularly done in cell-based phenotypic assays, but it is often unclear how to identify potential molecular targets in these experiments [9-11]. Understanding how cells respond when specific interactions are disrupted is not only essential for target identification but also for developing therapies that might restore perturbed disease networks to their native states.

Compound-centric computational approaches are now commonly applied to predict drug–target interactions by leveraging existing data. However, many of these methods extrapolate from known chemistry, structural homology, and/or functionally related compounds, and excel in target prediction only when the query compound is chemically or functionally similar to known drugs [12-17]. Other structure-based methods such as molecular docking are able to evaluate novel chemistries, but are limited by the availability of protein structures [18-20], inadequate scoring functions, and excessive computing times, which render structure-based methods ill-suited for genome-wide virtual screening [21].

More recently, a new paradigm for predicting molecular interactions using cellular gene expression profiles has emerged [22-24]. Previous work has shown that distinct inhibitors of the same protein target produce similar transcriptional responses [25]. Related studies have predicted secondary pathways affected by known inhibitors by identifying genes that, when null-mutated, diminish the inhibitory expression signature of drug-treated cells [26]. When no target information is available for the compound in question or related compounds, alternate approaches have mapped drug-induced differential gene expression levels onto known protein interaction network topologies and prioritized potential targets by identifying highly perturbed subnetworks [27-29]. These studies predicted roughly 20% of known targets within the top 100 ranked genes (see Methods for details), but did not predict or validate any previously unknown interactions.

The NIH’s Library of Integrated Cellular Signatures (LINCS) project presents an opportunity to leverage gene expression signatures from other types of cellular perturbations for the purpose of drug-target interaction prediction. Specifically, the LINCS L1000 dataset contains cellular mRNA signatures from treatments with 20,000+ small molecules and 20,000+ gene over-expression (cDNA) or knockdown (sh-RNA) experiments. Based on the hypothesis that drugs which inhibit their target(s) should yield similar network-level effects to silencing the target gene(s) (Figure 1a), we calculated correlations between the expression signatures of thousands of small molecule treatments and gene knockdowns in the same cells. We used the strength of these correlations to rank potential targets for a validation set of 29 FDA-approved drugs tested in the seven most abundant LINCS cell lines. We evaluate both direct signature correlations between drug treatments and knockdowns of their potential targets, as well as indirect signature correlations with knockdowns of proteins up/down-stream of potential targets. We combined these correlation features with additional gene annotation, protein interaction and cell-specific features in a supervised learning framework and use Random Forest (RF) [30, 31] to predict each drug’s target, achieving a top 100 target prediction accuracy of 55%, which we show is due primarily to our novel correlation features.

**Figure 1.**
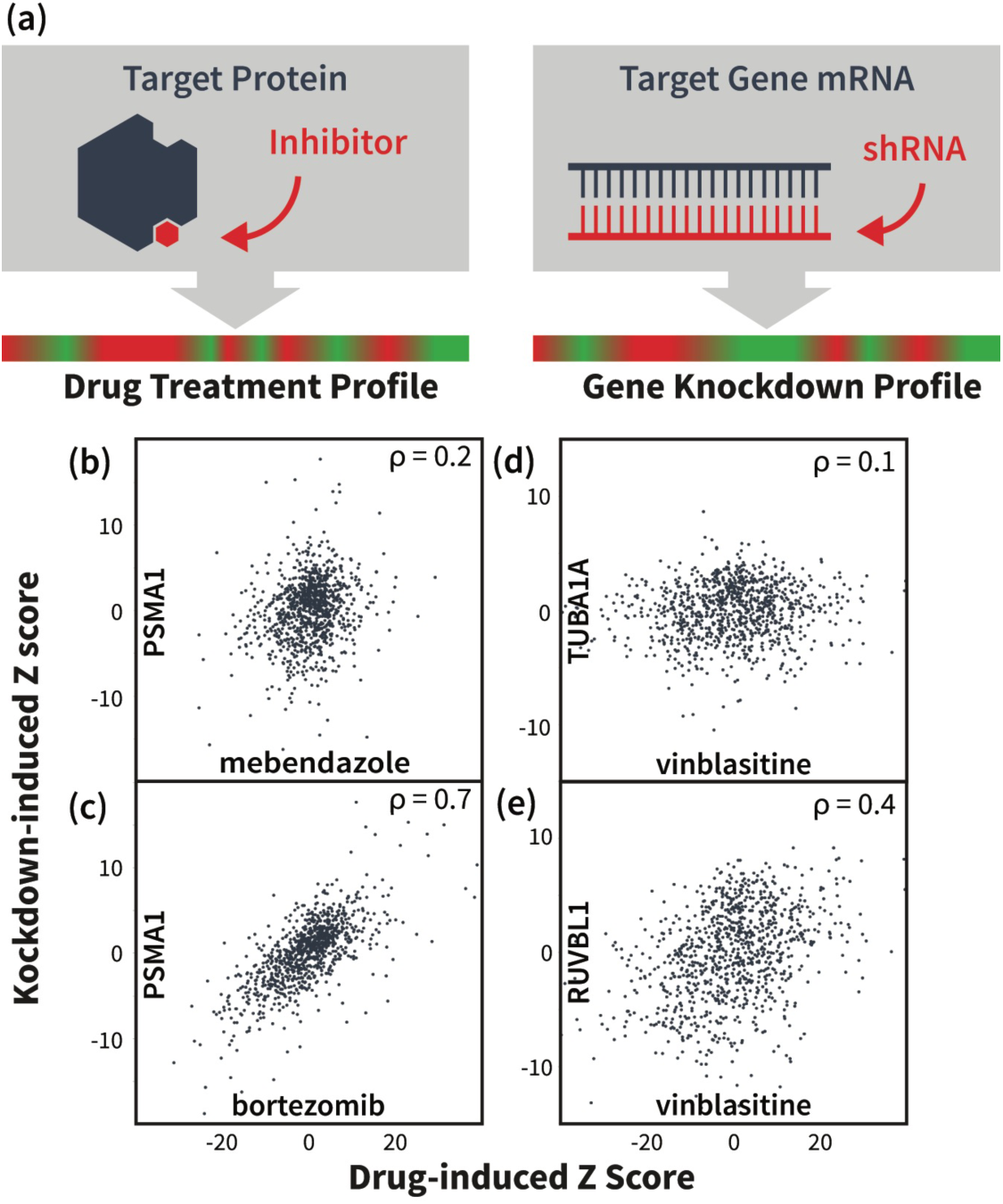
Drug and gene knockdown induced mRNA expression profile correlations reveal drug-target interactions. (a) Illustration of our main hypothesis: we expect a drug-induced mRNA signature to correlate with the knockdown signature of the drug’s target gene and/or genes on the same pathway(s). (b,c) mRNA signature from knockdown of proteasome gene PSMA1 does not significantly correlate with signature induced by tubulin-binding drug mebendazole, but shows strong correlation with signature from proteasome inhibitor bortezomib. Data points represent differential expression levels (Z-scores) the 978 landmark genes measured in the LINCS L1000 experiments. (d,e) Signature from tubulin-binding drug vinblastine shows little signature correlation with knockdown of its target TUBA1A, but instead correlates with the knockdown of functionally related genes, such as RUVBL1.

Finally, to filter out false positives and further enrich our predictions, we used molecular docking to evaluate the structural compatibility of the RF-predicted compound–target pairs. This orthogonal analysis significantly improved our prediction accuracy on an expanded validation set of 152 FDA-approved drugs, obtaining top-10 and top-100 accuracies of 26% and 41%, respectively, more than double that of aforementioned previous methods. We applied our pipeline to 1680 small molecules profiled in LINCS and experimentally validated seven potential first-in-class inhibitors for high-impact cancer targets HRAS, KRAS, CHIP, and PDK1.

These novel inhibitor candidates validated our hypothesis that drug treatments and target knockdowns cause similar disruptions of cellular protein networks that produce directly correlating differential expression patterns. More interestingly, we discover that these correlations can occur for knockdowns of the drug’s actual protein target(s) and/or for genes up/downstream of the target(s). We refer to the latter as “indirect correlations”. Several aspects of our approach represent significant step forwards from previous work exploring expression correlations as a means of predicting molecular interactions [32, 33]. Primarily, we do not assume anything about the small molecule or its likely protein target/pathway and our evaluation of both direct and indirect correlations allow us to screen compounds at a much larger scale and with higher accuracy than has been done previously. Furthermore, to our knowledge, this is the first time that pathway connectivity is explicitly considered by indirect correlational effects between drugs and knockdowns of target interaction partners. This approach helps to visualize and quantify the impact of drugs at the cell level and significantly improves in the translational potential of gene expression data to various realms of chemical biology and medicine. Finally, we open source our predictions and methods, providing enriched sets of likely active compounds for hundreds of human targets and presenting a new avenue for identifying suitable (multi-) targets for novel chemistries and accelerating the discovery of chemical probes of protein function.

## Results

### Preliminary prediction of drug targets using expression profile correlation features

We constructed a validation set of 29 FDA-approved drugs that had been tested in at least seven LINCS cells lines, and whose known targets were among 2634 genes knocked down in the same cell lines. For these drugs, we ranked potential targets using the direct correlation between the drug-induced mRNA expression signature and the knockdown-induced signatures of potential targets (Figure 1b,c). For each cell line, the 2634 knockdown signatures were sorted by their Pearson correlation with the expression signature of the drug in that cell line. We used each gene’s lowest rank across all cell lines to produce a final ranking of potential targets for the given drug. Using this approach, we predicted known targets in the top 100 potential targets for 8/29 validation compounds (Table 1). Indirect correlations were evaluated by the fraction of a potential target’s known interaction partners (cf. BioGrid [34]) whose knockdown signatures correlated strongly with the drug-induced signature. Ranking by indirect correlations predicted the known target in the top 100 for 10 of our 29 validation compounds (Table 1). Interestingly, several of these compounds showed little correlation with the knockdown of their targets (Figure 1d,e), with only 3/10 targets correctly predicted using the direct correlation feature alone.

It is well known that expression profiles vary between cell types [35]. Thus, we constructed a cell selection feature to determine the most “active” cell line, defined as the cell line producing the lowest correlation between the drug-induced signature and the control signature. Ranking by direct correlations within the most active cell line for each drug predicted six known targets in the top 100 (Table 1). However, all six of these targets were already predicted by either direct or indirect correlations, strongly suggesting that scanning for the optimal correlation across all cell lines is a better strategy than trying to identify the most relevant cell type by apparent activity.

**Table 1.** Performance of target prediction using different features and methods on the 29 FDA-approved drugs tested in 7 cell lines. DIR: direct correlation feature; IND: indirect correlation feature; CS: cell selection feature; MAX: maximum differential expression feature; MEAN: mean differential expression feature; LR: logistic regression; RF: random forest. Values are for the ranking of the top known target for each drug.

| Drug | Random | DIR | IND | CS | MAX | MEAN | LR | RF |
| --- | --- | --- | --- | --- | --- | --- | --- | --- |
| vinorelbine | 310 | 126 | 128 | 1318 | 1690 | 425 | 28 | 88 |
| dexamethasone | 1498 | 1891 | 284 | 943 | 315 | 1143 | 757 | 157 |
| dasatinib | 2325 | 1009 | 94 | 222 | 290 | 2621 | 182 | 532 |
| vincristine | 1979 | 473 | 439 | 386 | 2231 | 2196 | 456 | 37 |
| mycophenolate-mofetil | 564 | 1100 | 1263 | 2986 | 100 | 301 | 3064 | 3086 |
| amlodipine | 995 | 1338 | 2439 | 1801 | 1875 | 974 | 3037 | 650 |
| lovastatin | 1712 | 72 | 811 | 2078 | 1124 | 1068 | 1334 | 55 |
| clobetasol | 2194 | 820 | 21 | 157 | 74 | 15 | 38 | 65 |
| calcitriol | 2514 | 1059 | 2938 | 221 | 125 | 1814 | 1299 | 252 |
| flutamide | 919 | 2604 | 69 | 2806 | 463 | 298 | 702 | 647 |
| prednisolone | 2382 | 1439 | 206 | 787 | 402 | 1068 | 257 | 23 |
| nifedipine | 940 | 1225 | 1465 | 1285 | 88 | 322 | 3037 | 2249 |
| vemurafenib | 1042 | 1 | 82 | 1 | 1149 | 1403 | 22 | 2 |
| glibenclamide | 29 | 1415 | 2028 | 409 | 1059 | 740 | 1300 | 366 |
| digoxin | 2376 | 73 | 1470 | 118 | 828 | 567 | 732 | 44 |
| bortezomib | 1882 | 1 | 1 | 2 | 2546 | 2513 | 24 | 5 |
| vinblastine | 1612 | 515 | 56 | 100 | 224 | 377 | 38 | 2 |
| digitoxin | 573 | 89 | 430 | 216 | 521 | 653 | 79 | 50 |
| losartan | 645 | 489 | 988 | 770 | 636 | 31 | 735 | 1931 |
| pitavastatin | 1855 | 1976 | 1036 | 1117 | 90 | 527 | 1632 | 373 |
| digoxin | 69 | 521 | 776 | 194 | 127 | 559 | 208 | 64 |
| hydrocortisone | 303 | 312 | 72 | 58 | 93 | 122 | 29 | 17 |
| paclitaxel | 2299 | 74 | 121 | 47 | 371 | 1862 | 79 | 19 |
| lovastatin | 988 | 1 | 735 | 1587 | 1698 | 1484 | 128 | 100 |
| irinotecan | 1742 | 1023 | 20 | 236 | 128 | 1886 | 46 | 160 |
| vincristine | 1394 | 96 | 74 | 17 | 1272 | 69 | 28 | 9 |
| vinblastine | 1359 | 490 | 75 | 1383 | 373 | 1735 | 35 | 2 |
| raloxifene | 2080 | 2883 | 1818 | 1172 | 1064 | 479 | 1114 | 2520 |
| digoxin | 1005 | 102 | 1066 | 112 | 2096 | 2027 | 252 | 167 |
| Mean Ranking | 1365 | 800.6 | 724.3 | 776.9 | 794.9 | 1009.6 | 712.8 | 471.4 |
| Top 100 | 2 | 8 | 10 | 6 | 5 | 3 | 11 | 16 |

Finally, to incorporate the findings of previous studies that suggest that drug treatments often up/down regulate the expression of their target’s interaction partners [27-29], we constructed two features to report directly on the drug-induced differential expression of potential targets’ interaction partners. These features compute the maximum and the mean differential expression levels of potential targets’ interaction partners in the drug-induced expression profile. The lowest rank of each potential target across all cell lines is used in a final ranking. Though neither expression feature produces top 100 accuracies better than those of our correlation features, maximum differential expression identifies three new targets that were not identified using any of the previous features (Table 1).

#### Combining individual features using random forest (RF)

While each of the features in Table 1 performed better than random, combining them further improved results. Using Leave-One-Out Cross Validation (LOOCV) for each drug, logistic regression [31] correctly identified known targets in the top 100 predictions for 11 out of 29 drugs and improved the average known target ranking of all drugs (Table 1). However, logistic regression assumes that features are independent, which is not the case for our dataset given the complexity and density of cellular protein interaction networks. Hence, we used RF, which is able to learn more sophisticated decision boundaries [36]. Following the same LOOCV procedure, the RF classifier led to much better results than the baseline logistic regression, correctly finding the target in the top 100 for 16 out of 29 drugs (55%) (Table 1). Without further training, we tested the RF approach on the remaining 123 FDA-approved drugs that had been profiled in 4, 5, and 6 different LINCS cell lines, and whose known targets were among 3104 genes knocked down in the same cells. We predicted known targets for 32 drugs (26%) in the top 100 (Additional File 1), an encouraging result given the relatively small size of the training set and the expected decline in accuracy as the number of cell lines decreases (Table 2).

**Table 2.** Performance of two random forest models on validation set of 152 FDA-approved drugs as a function of cells tested. The number of drugs with targets ranked in top 100/50 are shown for the “on-the-fly” and “two-level” RF classification models. Results are divided into subsets of drugs profiled in different numbers of cell lines. Note that the success rate for RF is significant with p < 10^-6^ based on randomization tests (Figure S1).

| # of Cells | All | 7 | 6 | 5 | 4 |
| --- | --- | --- | --- | --- | --- |
| # of Drugs | 152 | 29 | 30 | 42 | 51 |
| On-the-fly |  |  |  |  |  |
| Top 100 | 58 | 13 | 15 | 16 | 14 |
| Top 50 | 42 | 10 | 10 | 12 | 10 |
| Top 100% | 38% | 45% | 50% | 38% | 27% |
| Top 50% | 28% | 34% | 33% | 29% | 20% |
| Two-level |  |  |  |  |  |
| Top 100 | 63 | 14 | 15 | 22 | 13 |
| Top 50 | 54 | 12 | 14 | 20 | 8 |
| Top 100% | 41% | 48% | 50% | 52% | 25% |
| Top 50% | 36% | 41% | 47% | 48% | 16% |

Re-training on the full set of 152 drugs and validating using LOOCV, we tested two alternative RF models: “on-the-fly”, which learns drug-specific classifiers trained on the set of drugs profiled in the same cell types, and “two-level”, which learns a single classifier trained on experiments from all training drugs (see Methods). The performances of both methods as a function of the number of cell lines profiled are summarized in Table 2. On-the-fly RF correctly ranked the targets of 58 out of 152 drugs in the top 100 (38%), with 42 of them in top 50 (28%). Two-level RF produced better enrichment, correctly predicting targets for 63 drugs in the top 100 (41%), and for 54 drugs in the top 50 (36%). In sharp contrast, random rankings (based on 20000 permutations) leads to only 7% of drugs with targets in the 100, indicating that both our training/testing and LOOCV results are extremely significant (Figure S1). It is also noteworthy that the top-100 accuracy of the two-level RF analysis increases to 50% if we only consider drugs treated in 5 or more cell lines.]

#### Gene ontology analysis of protein targets

Next, we analyzed in what context our Random Forest analysis was most successful. To do this, we divided the 152 drugs in our training data into “successful” predictions (the 63 drugs for which the correct target was ranked in the top 100), and “unsuccessful” predictions. We also divided the known targets into those that were correctly predicted and those that were not. We considered several different ways to characterize small molecules including molecular weight, solubility, and hydrophobicity, but none of these seemed to significantly correlate with our “successful” and “unsuccessful” classifications. Next, we used gene ontology to test for enrichment of “successful” and “unsuccessful” targets. Interestingly, we found that “successful” targets were significantly associated with intracellular categories, while the “unsuccessful” targets were mostly associated with transmembrane and extracellular categories (Table S1).

Based on this result we further incorporated cellular component as a feature in our two-level RF. We encode this feature by assigning 1 to the intracellular genes and -1 to the extracellular ones. We ran the two-level random forest with this additional feature included and demonstrated that the cellular component increases the number of top 100 genes to 66 and top 50 genes to 55.

#### Structural enrichment of genomic predictions

Figures 1d,e show that the gene regulatory effects of TUBA1A inhibition by the drug vinblastine manifest primarily as indirect correlations with knockdowns of the target’s interaction partners, such as RUVBL1, rather than via direct correlation with knockdown of the target. Such cases reflect the intrinsic connectivity of cellular signaling networks, which sometimes produce gene expression correlations that are ambiguous with respect to which of the interacting proteins in the affected pathway is the drug’s actual target. Our pipeline eliminates some of these false positives using an orthogonal structure-based docking scheme that, although limited to targets with known structure, allows us to significantly improve our prediction accuracy. After performing RF classification on the validation set, we mined the Protein Data Bank (PDB) [37] to generated structural models of the potential targets for our 63 “hits” - drugs for which we correctly identified the known target in the top 100. We selected one or more representative crystal structures for each potential target gene, optimizing for sequence coverage and structural resolution (see Supplemental Methods). We then docked hits to their top 100 potential targets and ranked using a prospectively validated pipeline [38-41].

On average, crystal structures were available for 69 out of the top 100 potential targets for each compound, and structures of known targets were available for 53 of the 63 hits. In order to avoid redocking into cocrystals of our hits, we made sure to exclude from our analysis all crystal structures containing these 53 ligands, ensuring that our results would not depend on prior knowledge of interaction partners or binding modes. As shown in Figure 2, molecular docking scores improved the re-ranking of the known target for 40 of the 53 drugs, with a mean and median improvement of 13 and 9, respectively. Based on genomic data alone, the known target was ranked in the top 10 for 40% of the 63 hits. After structural re-ranking, 65% had their known targets in the top 10 candidates, and this value improved to 75% in the subset of 53 drugs with known target structures. These results demonstrate the orthogonality of the genomic and structural screens, showing that molecular docking can efficiently screen false positives in our gene expression-based predictions.

**Figure 2.**
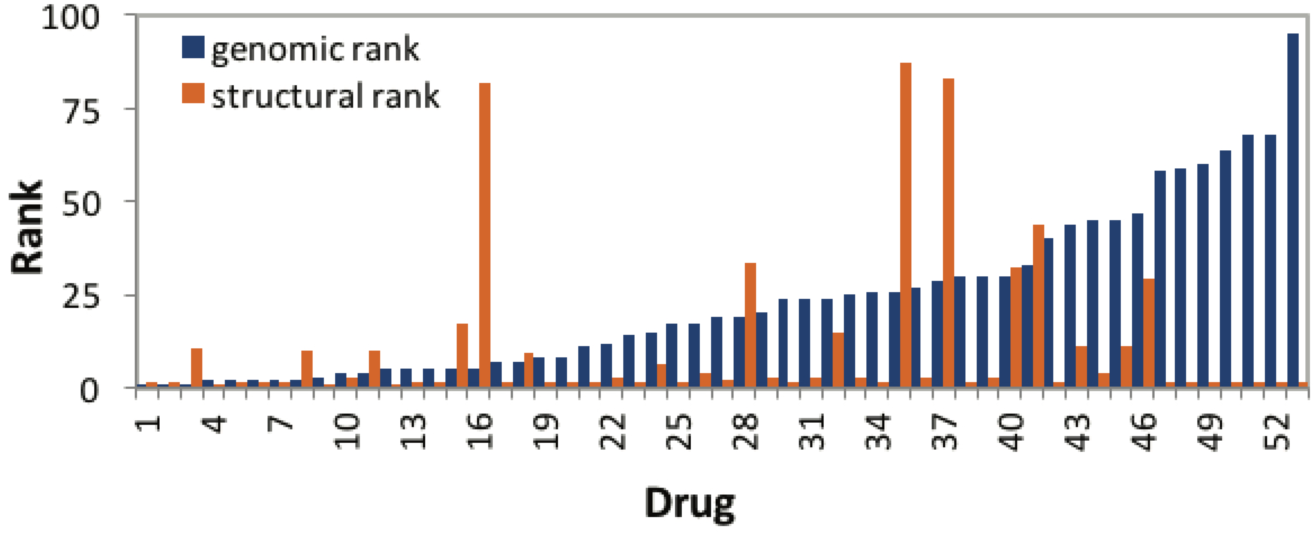
Structural enrichment of genomic target predictions. Predicted ranking (lower is better) of the highest-ranking known target for the 53 hits in our validation set with known target structures. Percentile rankings are shown following RF analysis (blue), and following structural re-ranking (orange). Drug names/IDs are listed in Additional File 2.

#### Identifying new interactions in the LINCS dataset

After validating our approach on known drug targets, we applied our pipeline to a test set of 1680 small molecules and 3333 gene knockdowns and predicted several novel interactions. We applied our pipeline (Figure 3) in both compound-centric (target prediction) and target-centric (virtual screening) contexts, in each case producing a final, enriched subset of roughly 10 predictions (either compounds or targets) that we tested experimentally. In compound-centric analyses, we performing molecular docking on the available structures of the input compound’s top-100 RF-predicted targets. In target-centric analyses, we ran the RF on our full test set, identified compounds for which the input protein is ranked in the top 100 potential targets, and then docked these candidate inhibitors to the target. In both applications, we analyzed the final docking score distributions and applied a 50% cutoff threshold to identify highly enriched compound/target hits. Structural analysis further facilitated visual validation of the docking models of predicted hits, thereby minimizing false positives.

**Figure 3.**
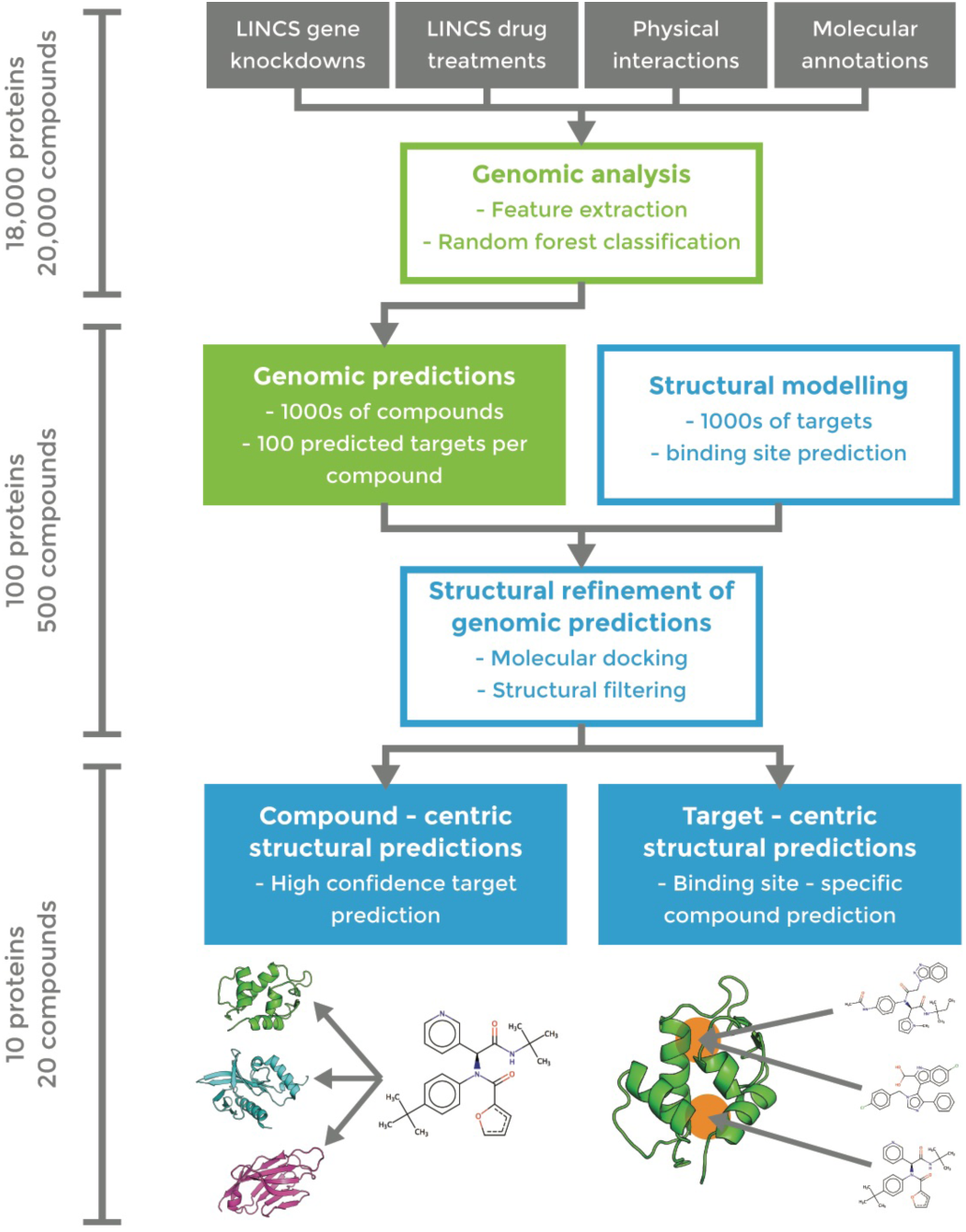
Workflow of combined genomic (green) and structural (blue) pipeline for drug-target interaction prediction. Approximate numbers of proteins/compounds in each phase are indicated on the left.

#### Target-centric prediction of novel RAS inhibitors

Our first application consisted in identifying novel binders of the high-impact and historically “undruggable” RAS-family oncoproteins. HRAS and KRAS are among the most frequently mutated genes in human cancers [42, 43]. However, despite the extensive structural data available and tremendous efforts to target them with small-molecule therapeutics, as of yet no RAS-targeting drug candidates have shown success in clinical trials [44-46].

Among the 1680 compounds in our test set, 84 and 156 were predicted (within the top-100) to target KRAS and HRAS, respectively. These compounds produced mRNA perturbation signatures that correlated strongly with knockdowns of KRAS (Figure 4a), and HRAS (Figure 4b). Of note, differential expression of genes functionally related to K/HRAS, i.e. FGFR4, FGFR2, FRS1, inform on novel regulatory phenotypes responding to both compound inhibition and gene knock out. We docked predicted compounds to our representative structures of KRAS (PDB ID: 4DSO [44]) and HRAS (PDB ID: 4G0N [47]) (Figure 4c,d). RF ranking and docking score distributions were compared to select compounds from our enriched datasets that were both commercially available and moderately priced. Docking models of promising candidates were also examined visually such that to reject models with unmatched hydrogen bonds [48] and select those that showed suitable mechanisms of action (see, e.g., Figure 4c,d). We purchased six potential HRAS inhibitors and five potential KRAS inhibitors for experimental validation (Table S2).

**Figure 4.**
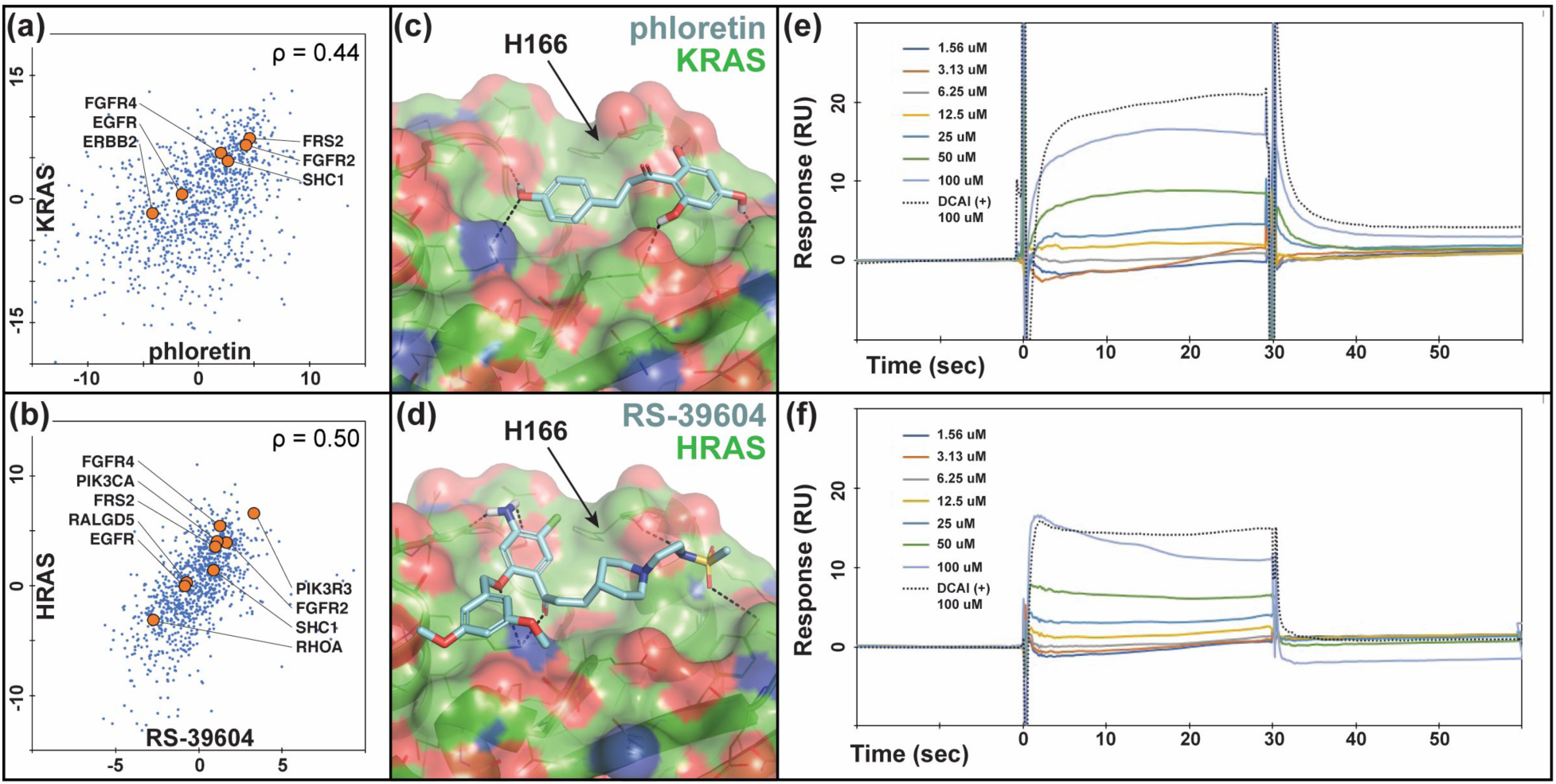
HRAS/KRAS inhibitors predicted based on direct correlations and docked poses show direct binding in SPR assays. Differential gene expression profiles of (a) Phloretin and (b) RS-39604 cell treatments and KRAS and HRAS knockdown experiments, respectively. Several functionally related genes listed in BioGrid [34] are indicated to demonstrate the relevance of these profiles as suggestive of direct drug-target interactions. Models of (c) phloretin and (d) RS-39604 bound to an allosteric site on the KRAS and HRAS catalytic domains, respectively. (e) SPR titration response curves for (e) phloretin and (f) RS-39604 binding to KRAS and HRAS, respectively, compared to DCAI positive control.

Our SPR assay measured direct binding of predicted inhibitors to AviTagged HRAS and KRAS. Initial 100 µM screens showed binding response for compounds RS-3906 against HRAS and phloretin against KRAS, and subsequent titrations confirmed binding at µM concentrations (Figure 4e,f), comparable to the DCAI positive control [44].

#### Target-centric prediction of novel CHIP inhibitors

Next, we targetted STUB1, also known as CHIP (the carboxy-terminus of Hsc70 interacting protein), an E3 ubiquitin ligase that manages the turnover of over 60 cellular substrates [49], which to our knowledge lacks specific inhibitors. CHIP interacts with the Hsp70 and Hsp90 molecular chaperones via its TPR motif, which recruits protein substrates and catalyzes their ubiquitination. Thus, treatment with small molecules that inhibit CHIP may prove valuable for pathologies where substrates are prematurely destroyed by the ubiquitin-proteasome system [50].

The screening of the 1680 LINCS small molecules profiled in at least four cell lines predicted 104 compounds with CHIP among the top 100 targets. We docked these molecules to our representative structure of the TPR domain of CHIP (PDB ID: 2C2L [51]), for which we had an available fluorescence polarization (FP) assay. The RF ranking and docking score distributions were compared to select compounds highly enriched in one or both scoring metrics. We next visually examined the docking models of top ranking/scoring hits to select those that show suitable mechanisms of action, and purchased six compounds for testing (Table S3). In parallel, we performed a pharmacophore-based virtual screen of the ZINC database [52] using the *ZincPharmer* [41] server, followed by the same structural optimization [38-41] performed on the LINCS compounds. We purchased seven of the resulting ZINC compounds for parallel testing.

Our FP assay measured competition with a natural peptide substrate for the CHIP TPR domain. We found that four (out of six) of our LINCS compounds reliably reduced substrate binding (Figure 5a,b), while three (out of seven) ZINC compounds did so to a modest degree (Figure S2). The two strongest binders were LINCS compounds 2.1 and 2.2. A functional assay also verified that 2.1 and 2.2 prevented substrate ubiquitination and CHIP autoubiquitination (Figure 5c,d, Figure S3). Compounds 2.1 and 2.2 also prevented ubiquitination of an alternate substrate that was tested subsequently (Figure S4). Importantly, the predicted binding modes of these two compounds did not match the pharmacophore model of the TPR-HSP90 interaction [51], which was used to screen the ZINC database (Figure S5). The latter emphasizes the power of our approach to identify novel compounds and mechanisms of action to targets without known inhibitors.

**Figure 5.**
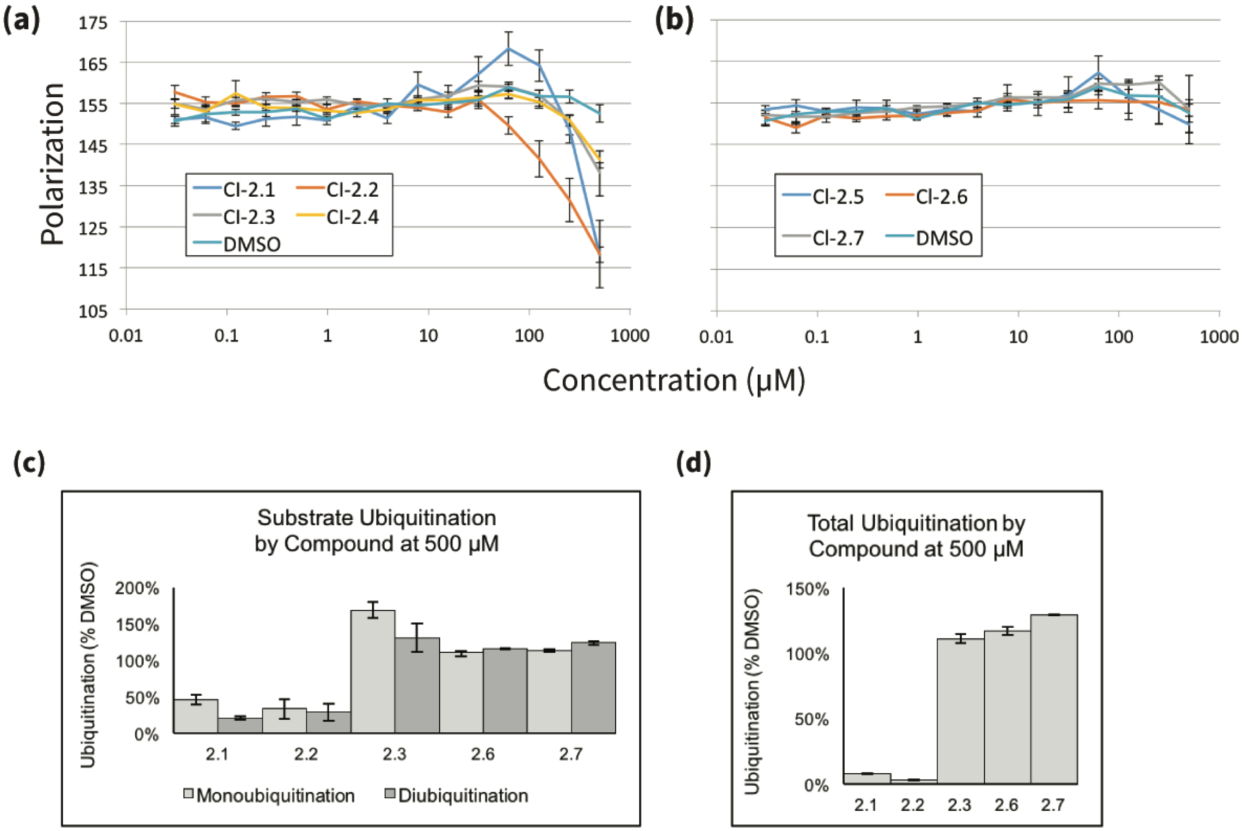
Predicted inhibitors show direct binding to and functional inhibition of CHIP. (a,b) Predicted CHIP inhibitors disrupt binding to chaperone peptide by fluorescence polarization. High ranked (a) and low ranked (b) compounds were tested for the ability to compete with a known TPR ligand (5-FAM-GSGPTIEEVD, 0.1 µM) for binding to CHIP (0.5 µM). Results are the average and standard error of the mean of two experiments each performed in triplicate. (c,d) CHIP inhibitors prevent ubiquitination by CHIP in vitro. (c) Quantification of substrate ubiquitination by CHIP from Anti-GST western blot experiments with tested compounds at 500µM, blotted as in Figure S3a and normalized to DMSO treated control (2.1, 2.2: N=4; all other compounds: N=2). (d) Quantification of total ubiquitination by CHIP from Anti-GST western blot experiments with tested compounds at 500µM, blotted as in Figure S3b and normalized to ubiquitination by a DMSO treated control (all compounds: N=2).

Contrary to the RAS compounds that were identified based on direct correlations between compound treatments and RAS knockdowns (Figure 4a,b), CHIP hits show almost no direct correlation (*ρ*_2.1;_ = 0.15, *ρ*_2.2_= 0.02), but were predicted based on indirect correlations with CHIP interaction partners. Figure 6 shows the correlating differential gene expression profiles for compound 2.1 and knockdowns of the CHIP interaction partners UbcH5 and HSP90, which, along with CHIP, were also predicted as potential targets by the RF classifier. However, structural screening ruled out these two partners as potential targets because of a lack of favorable binding modes.

**Figure 6.**
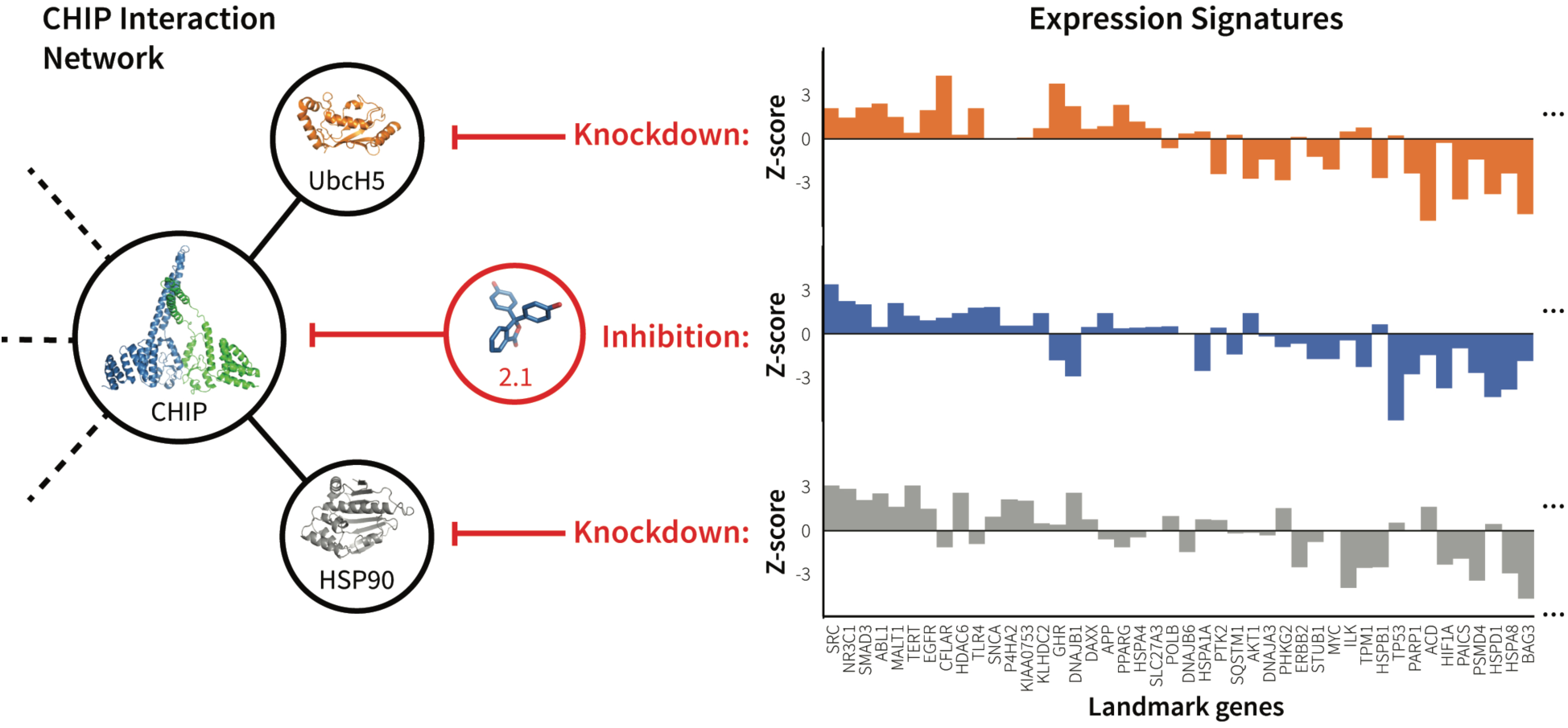
mRNA expression signature of CHIP inhibitor 2.1 correlates with knockdown of CHIP interacting partners. The figure illustrates the correlation between the mRNA expression profile signatures produced by treating cells with 2.1 and by knocking down CHIP interaction partners UbcH5 and HSP90. These three perturbations have similar network effects (left), as illustrated by their resulting differential expression signatures (right). For clarity, expression signatures show only the subset of LINCS landmark genes that are functionally related to CHIP according to BioGRID [34].

#### Compound-centric prediction of a novel target for the drug Wortmannin

We demonstrated a compound-centric application of our pipeline by analyzing Wortmannin, a selective PI3K covalent inhibitor and commonly used cell biological tool. DrugBank [53] lists four known human targets of Wortmannin: PIK3CG, PLK1, PIK3R1, and PIK3CA. Of the 100 targets predicted for Wortmannin, the PDB contained structures for 75, which we used to re-rank these potential targets. Only one known kinase target of Wortmannin, PIK3CA, was detected, and ranked 5^th^. Our pipeline also ranked 2^nd^ the human kinase PDPK1 (PDK1). Although PDK1 is a downstream signaling partner of PI3Ks [54], there is no prior evidence of a direct Wortmannin-PDK1 interaction in the literature. Nevertheless, both the strong direct correlation of wortmannin with the PDK1 knockdown (Figure 7a), and the native-like binding mode predicted by our pipeline (Figure 7b) suggested a possible interaction.

**Figure 7.**
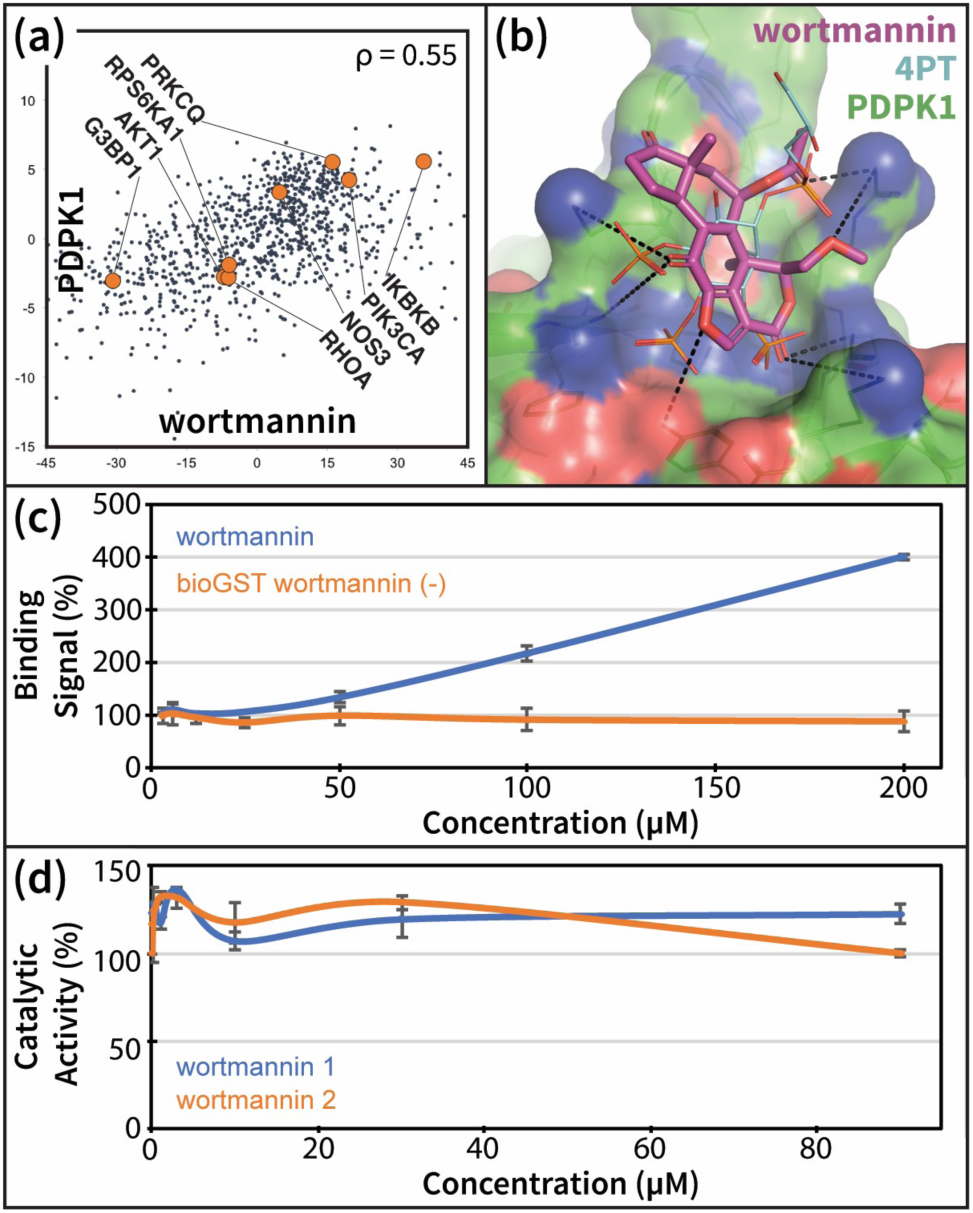
Wortmannin promotes PDK1 – PIP3 binding in vitro. (a) Wortmannin treatment and PDK1 knockdown experiments produce directly correlating differential gene expression profiles. Several functionally related genes listed in BioGrid [34] are indicated to demonstrate the relevance of these profiles as suggestive of direct drug-target interactions. (b) Model of wortmannin bound to the PH domain of PDK1, compared to known ligand 4PT (PDB ID: 1W1G [55]). (c) Alphascreen PDK1-PIP3 interaction-displacement assay results for increasing concentrations of wortmannin. Error bars represent the standard error on the mean from two parallel runs. (d) Effect of wortmannin on the in-vitro phosphorylation of the substrate T308tide by the isolated catalytic domain of PDK1. The two lines are from two replicates of the activity assay, with error bars representing the standard error on the mean from two parallel runs for each replicate.

We experimentally tested this interaction using an alphascreen PDK1 interaction-displacement assay. Since we predicted that Wortmannin binds to the PH domain of PDK1 (Figure 7b), we measured the effect of increasing Wortmannin concentrations on the interaction of PDK1 with the second messenger PIP3. We found that Wortmannin specifically increased PDK1-PIP3 interaction, relative to control (Figure 7c). Given that PIP3-mediated recruitment of PDK1 to the membrane is thought to play an important regulatory role in the activity of the enzyme [56, 57], a disruptive increase in PDK1-PIP3 interaction following treatment with Wortmannin supports our prediction.

#### Comparison to existing target prediction methods

For completeness, we compared results for our 63 hits from the validation set to those produced by available structure and ligand-based methods. HTDocking (HTD) [58] is a structure-based target prediction method that docks and scores the input compound against a manually curated set of 607 human protein structures. For comparison, in our analysis we were able to extract high-quality domain structures for 1245 (40%) of the 3104 potential gene targets. PharmMapper (PHM) [59] is a ligand-based approach that screens the input compound against pharmacophore models generated from publicly available bound drug-target cocrystal structures of 459 human proteins, and then ranks potential targets by the degree to which the input compound matches the binding mode of the cocrystalized ligands. The scope of HTD is limited by the availability of the target structure, while PHM is limited by chemical and structural similarity of active ligands.

HTD and PHM rankings for known targets are shown in Table 3, and complete results are shown in Additional File 3. Our combined genomics-structure method outperforms the structure-based HTD server (average ranking of the known target is 13 for our method vs. 50 for the HTD server). This suggests that limiting the structural screening to our genomic hits allowed us to predict targets with higher accuracy than docking alone. Results when using the PHM server are on average similar to ours. However, PHM relies on the availability of ligand-bound crystal structures, which in practice makes this class of methods more suitable for drug repurposing than assessing new chemistries or targets.

**Table 3.** Comparison of our pipeline to existing drug-target prediction methods. The average ranking of the highest ranked known target is listed for all 63 validation ‘hits’, for the subset of 53 validation hits with known target structures, and for our seven predicted interactions. ‘Structures available’ indicates the average number of top-100 potential targets with available crystal structures for the compound set. Rankings are compared between the initial random-forest genomic ranking, the structural re-ranking of the top 100 RF predicted targets, the HTDocking server (HTD), and the PharmMapper server (PHM).

|  | **Structures available** | **Genomic Rank** | **Structural Re-rank** | **HTD** | **PHM** |
| --- | --- | --- | --- | --- | --- |
| **All hits (n=63)** | 69 | 22 | 24 | 56 | 23 |
| **Hits w/ known target structures (n=53)** | 71 | 23 | 13 | 50 | 12 |
| **New predictions (n=7)** | 73 | 28 | 31 | n/a | n/a |

With regards to new validated interactions, alternative approaches failed to predict the interactions with HRAS, KRAS, and CHIP that were verified by our assays. However, a Wortmannin-PDK1 interaction was predicted at the catalytic site by HTD, ranked 540^th^, and by PHM, ranked 56^th^. Although we cannot rule out a possible kinase domain interaction, a catalytic activity assay showed that Wortmannin had no measureable effect on the in vitro phosphorylation of the substrate T308tide by the isolated catalytic domain of PDK1 (Figure 7d).

## Discussion

Delineating the role of small molecules in perturbing cellular interaction networks in normal and disease states is an important step towards identifying new therapeutic targets and chemistries for drug development. To advance on this goal, we developed a novel target prediction method based on the hypothesis that drugs that inhibit a given protein should have similar network-level effects to silencing the inhibited gene and/or its up/downstream partners. Using gene expression profiles from knockdown and drug treatment experiments in multiple cell types from the LINCS L1000 dataset, we developed several correlation-based features and combined them in a random forest (RF) model to predict drug-target interactions.

On a validation set of 152 FDA-approved drugs we achieve top-100 target prediction accuracy more than double that of previous approaches that use differential expression alone [28, 29]. Consistent with our underlying hypothesis, the RF results highlight the importance of both direct expression signature correlations between drug treatment and knockdown of the gene target (Figure 1c, Figure 4a,b, Figure 7a) and indirect correlations between the drug and the target’s interacting partners (Figure 1e, Figure 6). Contrary to earlier work [27-29], our method is capable of predicting potential targets for any compound, even those unrelated to known drugs, and our predictions are open source and available for immediate download and testing (http://sb.cs.cmu.edu/Target2/). These include potential targets for 1680 LINCS small molecules from among 3000+ different human proteins.

Unlike most available ligand-based prediction methods [12-17], the accuracy of our approach does not rely on chemical similarity between compounds in the training/test sets. For instance, our screen against CHIP, a target with no known small molecule inhibitors, delivered four out of six binding compounds, whereas a parallel analysis using a state-of-the-art structure-based virtual screening [38, 60] yielded only two weak-binding compounds. Moreover, the predicted mechanisms of actions of the more potent LINCS compounds suggest novel interactions that were not prioritized by the ligand-based screen (Figure S5).

In contrast to other machine learning methods, our approach reveals important, human-interpretable insights into perturbation-response properties of cellular networks. Direct and indirect gene expression profile correlations inform on global regulatory responses triggered by small molecule cell treatments (see, e.g., Figures 4, 6, 7). Namely, our genomic screening not only identifies compounds targeting a given protein, but also highlight related genes that are affected by the chemical modulation of the target. This knowledge is bound to play an important role in the design of polypharmacological therapies.

The experimental validation of our predictions for HRAS, KRAS, CHIP and Wortmannin demonstrate the power of our combined genomic and structural pipeline in identifying novel targets and chemotypes. Our prospectively identified modulators are the first of their kind – in that they represent the results of a virtual target-screening process, rather than traditional high-throughput small molecule screening approaches.

Detailed analyses of our predictions suggest several avenues to improve enrichment. We established a clear correlation between the number of cell-types screened and the target prediction accuracy. We identified that a significant source of false positives are indirect correlations that while important to detect the true target, also tend to predict interacting partners as potential targets. Incorporating compound- or target-specific features are also likely to improve our results. For instance, we noticed that our prediction results were less accurate for membrane proteins, and incorporating a cellular localization feature into our RF model increased the number of top-100 hits in our validation set from 63 to 66.

In sum, our method represents a novel application of gene expression data for small molecule– protein interaction prediction, with structural analysis further enriching hits to an unprecedented level in our proteome-scale screens. The success of our proof-of-concept experiments opens the door for a compound-centric drug discovery pipeline that can leverage the relatively small fraction of potentially bioactive compounds that could be of interest for further investigation to become drugs [61]. Compared to alternative approaches, our method would be particularly suitable for scanning for targets of newly synthesized scaffolds. We are hopeful that our open source method and predictions might be useful to other labs around the world for identifying new drugs for key proteins involved in various diseases and for better understanding the impact of drug modulation of gene expression. Moreover, our approach represents a new framework for extracting robust correlations from intrinsically noisy gene expression data that reflect the underlying connectivity of the cellular interactome.

## Materials and Methods

### Data sources

A full description of the data used in our analysis can be found in the Supplemental Methods. Briefly, from the NIH LINCS library we extracted gene expression perturbations on 978 “landmark genes” from thousands of small molecule treatment and gene knockdown experiments in various cell lines. We then used ChEMBL [62], an open large-scale bioactivity database, to identify the LINCS compounds that were FDA approved and had known targets. To construct our validation set we selected the 152 FDA approved compounds that had been tested in at least four distinct LINCS cell lines, and whose known targets were knocked down in the same cell lines. Protein-protein interaction data used in feature construction was extracted from BioGRID [34] and HPRD [63], both of which contain curated sets of physical and genetic interactions. Protein cellular localization data used in feature construction was obtained from the Gene Ontology database [64].

### Extracting and integrating features from different data sources

The notation and symbols that we use in constructing and using the genomic features are described in Table S4 and Table S5. Feature construction is summarized below and is explained in detail in the Supplemental Methods.

#### Direct correlation

The first feature *f*_*cor*_, computes the correlation between the expression profiles resulting from a gene knockdown and treatment with the small molecule. Since we are considering multiple cells for each molecule/knockdown, the correlation feature for each molecule *d*, i.e.*f*_*cor*_ (*d*,·,·), has a dimension of |*T*_*d*_| × |*C*_*d*_|.

#### Indirect correlation

Information about protein interaction networks may be informative about additional knockdown experiments that we might expect to be correlated with the small molecule treatment profile. To construct a feature that can utilize this idea we did the following: for each molecule, protein, and cell line we computed *f*_*PC*_ (*d, g, c*), which encodes the fraction of the known binding partners of *g* (i.e. the proteins interacting with *g*) in the top *X* knockdown experiments correlated with this molecule/cell compared to what is expected based on the degree of that protein (the number of interaction partners - this corrects for hub proteins). We used *X* = 100 here, though 50 and 200 gave similar results. See Supplemental Methods for complete details.

#### Cell selection

While the correlation feature is computed for all cells, it is likely that most drugs are only active in certain cell types and not others. Since the ability to consider the cellular context is one of the major advantages of our method we added a feature to denote the impact a drug has on a cell line. For each drug/molecule *d* we compute a cell specific feature, *f*_*CS*=_(d,·), which measures the correlation between the response expression profile and the control (WT) experiments for that cell. We expect a smaller correlation if the drug/molecule is active in this cell, and a larger correlation if it is not.

#### Differential expression

In addition to determining the correlation-based rankings of interacting proteins, we also took their drug-induced differential expression into account. We constructed two features that summarize this information for each protein (see Supplemental Methods for details). These features either encode the average or the max (absolute value) expression level of the interaction partners of the potential target protein.

### Generating structural models for docking

In order to use molecular docking to enrich of our random forest predictions, we needed to generate structural models for the genes profiled in LINCS. The union of our top 100 target predictions for the 1680 small molecules profiled in LINCS in at least four cell lines consisted of 3333 unique human genes. We used a python script (available on github^1^) to mine the PDB for structures of these genes and then select representative crystal structures for each. When multiple structures were available, a representative subset of structures were chosen so as to maximize sequence coverage, minimize strucutral resolution, and account for structural heterogeneity. Full details of this procedure can be found in the Supplemental Methods.

### Docking procedure

Compounds were docked to representative structures of their predicted targets with smina [39], using default exhaustiveness and a 6 Å buffer to define the box around each potential binding site. Docked poses across predicted binding sites [65] on a given target were compared and the highest scoring pose of each compound was selected for further analyses [38-41] and comparison to other targets/compounds.

### Experimental assays

Full details on all experimental assays involving HRAS, KRAS, CHIP and PDK1 can be found in the Supplemental Methods.

## Additional Files

**Additional File 1** (csv). Results of testing our random forest classifier on the 123 FDA approved drugs profiled in 4-6 LINCS cell lines after having trained our model on the 29 FDA approved drugs profiled in all 7 LINCS cell lines. The rank of the highest-ranking known target for each compound is listed next to their LINCS ID. We achieve top-100 predictions for 32 drugs, a 26% success rate.

**Additional File 2** (csv). The names and LINCS IDs of the validation compounds shown in Figure 2.

**Additional File 3** (csv). Structural enrichment of random forest predictions for validation hits and comparison with existing methods. Table lists the 63 `hits’ from our validation drug set, including their names, LINCS ID and the number of top-100 predicted targets that had structures available in the PDB. The ranking of the known targets for each compound are shown after our genomic random forest target prediction (GEN), and after our structural re-ranking (STR), along with the percentile rankings produced by alternative target prediction methods HTDocking (HTD) and PharmMapper (PHM). STR, HTD, and PHM values of 100 indicate that the structure of the known target either is not known or was not included in the set of potential targets used by the method.

## Acknowledgements

This work was supported in part by the U.S. National Institutes of Health (grants 1U54HL127624 to Z.B.J. and R01GM097082, to C.J.C.) and by the U.S. National Science Foundation (grants DBI-1356505 to Z.B.J. and 1247842, to N.A.P.). We thank the Connectivity Map team at the Broad Institute for generation of the LINCS data set and query tools. The authors declare no competing financial interests. We are also grateful to Andy Stephen, Ph.D., the team lead of the RAS Biochemistry and Biophysics Group at Leidos Biomedical Research, Inc., for his help validating our HRAS/KRAS predictions. In this work, proteins were produced by Troy Taylor, Shelley Perkins, John-Paul M. Denson and William Gillette. Biacore binding experiments were performed Patrick Alexander and Andrew Stephen, at the Frederick National Laboratory for Cancer Research, Frederick MD.

## TOC Graphic

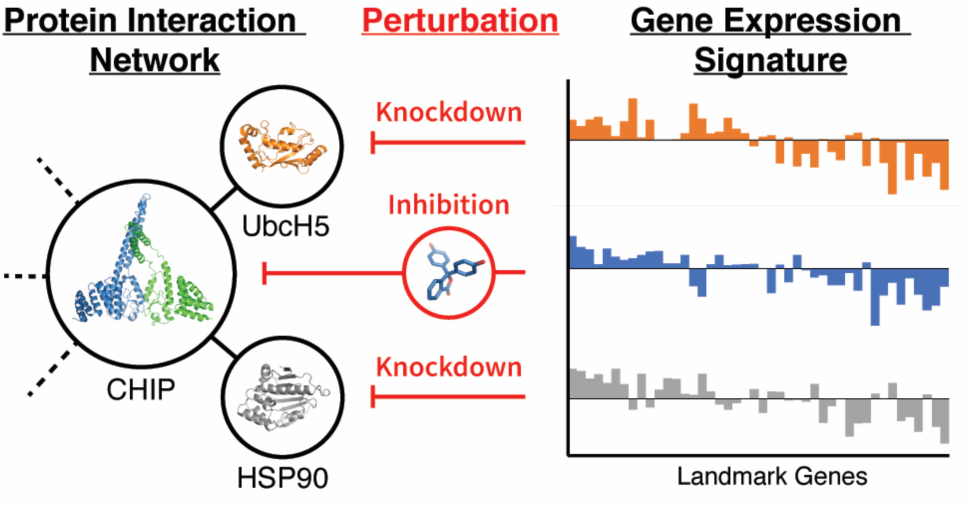

